# Coexistence between similar invaders: The case of two cosmopolitan exotic insects

**DOI:** 10.1101/2022.02.03.479030

**Authors:** Matthew B. Arnold, Michael Back, Michael Daniel Crowell, Nageen Farooq, Prashant Ghimire, Omon A. Obarein, Kyle E. Smart, Trixie Taucher, Erin VanderJeugdt, Kayla I. Perry, Douglas A. Landis, Christie A. Bahlai

**Affiliations:** Department of Biological Sciences, Kent State University, Kent, Ohio, United States of America, 44242; Department of Geography, Kent State University, Kent, Ohio, United States of America, 44242; Department of Geology, Kent State University, Kent, Ohio, United States of America, 44242; Department of Entomology, and Great Lakes Bioenergy Research Center, Michigan State University, East Lansing, Michigan, United States of America, 48824; Kellogg Biological Station, Michigan State University, Hickory Corners, Michigan, United States of America, 49060

**Keywords:** Biological control, *Coccinella septempunctata*, Coccinellidae, *Harmonia axyridis*, invasion, niche partitioning, coexistence

## Abstract

Biological invasions are usually examined in the context of their impacts on native species. However, few studies have examined the dynamics between invaders when multiple exotic species successfully coexist in a novel environment. Yet, long-term coexistence of now established exotic species has been observed in North American lady beetle communities. Exotic lady beetles *Harmonia axyridis* and *Coccinella septempunctata* were introduced for biological control in agricultural systems and have since become dominant species within these communities. In this study, we investigated coexistence via spatial and temporal niche partitioning among *H. axyridis* and *C. septempunctata* using a 31-year dataset from southwestern Michigan, USA. We found evidence of long-term coexistence through a combination of small-scale environmental, habitat, and seasonal mechanisms. Across years, *H. axyridis* and *C. septempunctata* experienced patterns of cyclical dominance likely related to yearly variation in temperature and precipitation. Within years, populations of *C. septempunctata* peaked early in the growing season at 550 degree days, while *H. axyridis* populations grew in the season until 1250 degree days, and continued to have high activity after this point. *Coccinella septempunctata* was generally most abundant in herbaceous crops, whereas *H. axyridis* did not display strong habitat preferences. These findings suggest that within this region *H. axyridis* has broader habitat and abiotic environmental preferences, while *C. septempunctata* thrives under more specific ecological conditions. These ecological differences have contributed to the continued coexistence among these two invaders. Understanding mechanisms that allow coexistence of dominant exotic species contributes to native biodiversity conservation management of invaded ecosystems.

**Open research statement:** Data are already published and publicly available, with those items properly cited in this submission. This submission uses novel code, which is provided, per our requirements, in an external repository to made available in perpetuity, and are available at https://github.com/ReproducibleQM/space_invader. Data sets utilized for this research (Landis 2020) are housed at EDI here: https://portal.edirepository.org/nis/mapbrowse?packageid=knb-lter-kbs.23.30 (doi:10.6073/pasta/f0776c1574808b08c484c1f7645a7357). Weather data was downloaded directly from the Kellogg Biological Station data repository (https://lter.kbs.msu.edu/datatables/7) and downloading the full record. An archival record of these data are available at https://portal.edirepository.org/nis/mapbrowse?packageid=knb-lter-kbs.2.107 (doi:10.6073/pasta/4c30523bae14c4340e4d9c90e72f90c4). Because both databases are ‘living’ and subject to update as data is collected, databases as used within this study are mirrored within the code repository as CSV files.

## Introduction

The establishment and spread of exotic species is a major driver of global environmental change, threatening native biodiversity, ecosystem services and function, and human well-being (Vitousek et al. 1996, Ricciardi 2007). Invasions of insect species have occurred on a global scale through intentional or unintentional introductions, resulting in substantial economic and ecological impacts (Kenis et al. 2009, Bradshaw et al. 2016). Ecological impacts of exotic insects affect native species directly and indirectly (Kenis et al. 2009, Vilà et al. 2011, Pyšek et al. 2020). For example, when exotic insects can successfully establish and spread outside their historical ranges, novel communities are formed where native and exotic species may interact directly through herbivory, predation, and parasitism (Liebhold et al. 1995, Boettner et al. 2000, Holway et al. 2002). Additionally, exotic species may cause indirect and/or cascading ecological impacts via various mechanisms such as exploitative and apparent competition, disease transmission, and alteration of habitat or food resources (Louda et al. 1997, Morin et al. 2007, Gandhi and Herms 2010, Klooster et al. 2018). As the frequency of invasions continues to increase globally (Lockwood et al. 2013, Seebens et al. 2017), research investigating novel interactions among native and exotic species is essential for assessing impacts of invaders as well as developing management strategies to conserve biodiversity.

Successful invasion of native communities by exotic species is context dependent and influenced by multiple factors such as abiotic environmental conditions, properties of the native community, and characteristics of the invading species (Blackburn et al. 2011, Lockwood et al. 2013). Because establishment success is influenced by local niche processes and interactions among species, hypotheses to explain the outcome of this stage in the invasion process draw on concepts within niche and coexistence theory (Shea and Chesson 2002, Godoy 2019). For example, the diversity-invasibility hypothesis (i.e. the biotic resistance hypothesis) predicts that more diverse native communities are more stable, and thus more resistant to the establishment of exotic species than less diverse communities (Jeschke 2014). This prediction is based on the premise that less diverse communities have more vacant niches available to invaders and a lower probability of occurrence of strong competitors or predators that could limit coexistence (Levine and D’Antonio 1999, Ricciardi et al. 2013). Long-term coexistence of exotic species within native communities is determined through stabilizing (niche differences) and equalizing (fitness differences) mechanisms wherein species vary in their environmental responses, resource acquisition, and/or competitive ability (Chesson 2000a, HilleRisLambers et al. 2012). Research has primarily focused on understanding interactions among native and exotic species to discern ecological impacts (e.g. Ricciardi et al. 2013). However, invasions have become widespread such that native communities are more commonly invaded by multiple exotic species which then directly or indirectly interact with each other to promote or inhibit coexistence.

Lady beetles (Coleoptera: Coccinellidae) are predatory insects that have been intentionally introduced for biological control in agricultural systems (Obrycki and Kring 1998, Koch 2003, Rondoni et al. 2021). This has led to the successful establishment and spread of several exotic lady beetles, including the Asian species *Harmonia axyridis* (Pallas) and the European species *Coccinella septempunctata* (Linnaeus) in North America. Both species are found in diverse habitats and primarily aphidophagous (Koch 2003, Hodek and Michaud 2008) but will feed on other arthropod prey and pollen if aphid resources are scarce (Berkvens et al. 2008, 2010, Evans 2009). The establishment and spread of *H. axyridis* and *C. septempunctata* have coincided with declines in native lady beetle species, while both invaders have been found to coexist (Alyokhin and Sewell 2004, Harmon et al. 2007, Roy et al. 2016). For example, *H. axyridis* and *C. septempunctata* were highly abundant within native lady beetle communities that were sampled over 24 years in southwestern Michigan, USA (Bahlai et al. 2015). Because these invaders have become dominant species within many native communities (Harmon et al. 2007, Bahlai et al. 2015, Gardiner et al. 2021), direct and indirect forms of competition are hypothesized as drivers of declines in native species (Pell et al. 2008).

Competitive interactions have been primarily investigated among native and exotic lady beetle species to assess mechanisms of decline and the impact of invasion (Pell et al. 2008, Roy et al. 2016, Rondoni et al. 2021). These exotic species share similar preferences in habitat and prey as some native lady beetles such that the degree of niche overlap with functionally similar invaders is hypothesized to drive competitive interactions (Snyder 2009). Apparent competition (Smith and Gardiner 2013) and intraguild predation (Gagnon et al. 2011, Thomas et al. 2013) have been observed among native and exotic species in the field, lending some support to this hypothesis. Evidence of exploitative competition also has been observed wherein competition from exotic species for shared prey has reduced weight gain and reproduction in some lady beetle species (Zaviezo et al. 2019) and shifted the habitat use patterns of native species from agricultural to natural environments such as forests (Evans 2004, Grez et al. 2013, Bahlai et al. 2015).

Less is known about competitive interactions among the two invaders *H. axyridis* and *C. septempunctata,* both of which are considered efficient competitors and capable of exploiting diverse habitats (Koch 2003, Hodek and Michaud 2008) but are known to coexist within similar environments. High levels of intraguild predation among larvae of *H. axyridis* and *C. septempunctata* have been observed in the laboratory (Snyder et al. 2004) and in the field (Gagnon et al. 2011). Outcomes of competitive interactions are influenced by the relative body size, mobility, age, and diet specificity of larvae as well as prey density (Hironori and Katsuhiro 1997, Yasuda et al. 2004). Larvae of *H. axyridis* tend to be larger and more aggressive than *C. septempunctata* (Yasuda et al. 2001, Ware and Majerus 2008), which may translate to an asymmetric competitive advantage. For example, larvae of *H. axyridis* were more successful at escaping attacks from *C. septempunctata* in laboratory pairwise experiments than vice versa, thereby reducing the survival of *C. septempunctata* larvae compared to *H. axyridis* larvae (Yasuda et al. 2001). Moreover, adults of *H. axyridis* found and consumed more aphids in the laboratory than other lady beetle species including *C. septempunctata* (Leppanen et al. 2012). Asymmetric competition in favor of *H. axyridis* suggests other forms of niche partitioning are likely facilitating coexistence of these invaders over time such as differential use of habitats and environmental preferences. For instance, colder minimum winter temperatures affect populations of both species, but effects are more strongly negative on *C. septempunctata* (Cheng et al. 2020). Increased overwintering survival of *H. axyridis* in very cold environments is often attributed to their behavior of hibernating in more temperate locations in buildings or under tree bark (Roy et al. 2016). High overwintering survival of *H. axyridis* following cold winters is predicted to lead to larger populations in spring and earlier reproduction than *C. septempunctata* (Raak-van der Berg et al. 2012), which could facilitate coexistence through alternating species dominance across years. Improved understanding of the mechanisms that allow coexistence and success of these dominant exotic species will inform biological control programs as well as biodiversity conservation management of invaded ecosystems.

To understand how *H. axyridis* and *C. septempunctata* have coexisted within native lady beetle communities over time, this study investigated environmental, habitat, and seasonal niche partitioning among these two dominant exotic species using a 31-year dataset from southwestern Michigan, USA. These long-term data allowed for the spatiotemporal analyses of abundances of both exotic species in nine plant habitats embedded within an agricultural landscape. Our goals were to evaluate environmental and ecological factors that may facilitate coexistence among these two species. We hypothesized that coexistence of these exotic species may occur via: 1) temporal niche partitioning with differing phenology among species; or 2) spatial niche partitioning with differing habitat preferences among species.

## Methods

Data for this study were collected at the Kellogg Biological Station Long-Term Ecological Research (KBS LTER) site located in southwest Michigan, USA. Our study focuses on data produced in the Main Cropping System Experiment (MCSE) at KBS LTER, a long-term agronomic experiment started in 1989, and the forest monitoring plots initiated in 1993 to document reference conditions adjacent to the MCSE site (Landis 2020). The experiment consists of an annual crop rotation (maize, soybean, wheat) maintained under four levels of management intensity, three perennial cropping systems (alfalfa/switchgrass, poplar tree plantation, and early successional vegetation maintained by yearly burnings), and three forest types (successional forest on reclaimed cropland, old growth deciduous fragments, and conifer plantations). In this study, we pooled data across management regimes by dominant plant community, totaling nine total habitats. Within each research plot, data were collected at five subsampling stations, with most measurements taken within the growing season (May to September of each year). Lady beetle populations are the focus of the insect survey, although several other taxa have also been recorded in more recent years (Colunga-Garcia et al. 1997, Colunga-Garcia and Gage 1998, Hermann et al. 2016).

The KBS LTER insect survey was established in 1989, soon after the arrival of *C. septempunctata* at the site (Maredia et al. 1992). *Coccinella septempunctata* is a large, primarily aphidophagous lady beetle believed to have been intentionally introduced to North America from Europe (Schaefer et al. 1987), but is now Holarctic in distribution (Hodek and Michaud 2008). By 1994, another invading lady beetle arrived at KBS LTER. Like *C. septempunctata, H. axyridis* is primarily aphidophagous and thrives in many habitat types with a now near-global distribution (Adriaens et al. 2008, Roy et al. 2016). Originally native to the Asian continent from northeastern China to Siberia (Roy et al. 2016), this species has been introduced as a biological control agent since the early 20^th^ century to North America and Europe (Roy et al. 2016, Sethuraman et al. 2018, Cheng et al. 2020). Although *C. septempunctata* is generally thought to be a European species, some sources note that the natural, or at least naturalized range of both *H. axyridis* and *C. septempunctata* have overlapped, and thus the two species have co-occurred in parts of China for quite some time (Cheng et al. 2020).

Insect surveys have been performed at the KBS LTER since 1989 using yellow sticky traps, placed at permanent sampling stations within the MSCE and forest sites. Yellow sticky traps were placed on T-posts, held at 1.2 m above the ground at each station during the growing season. Sample collection periods varied from 8-15 weeks each year depending upon crop management, environmental conditions, and labor availability (Bahlai et al. 2013, Bahlai, Colunga-Garcia, et al. 2015). Traps were inspected weekly, lady beetles captured were identified to species, and observations were recorded as the number of adults, by species, by date.

Insect data were examined at two different temporal resolutions. First, to match the typical sampling frequency during the growing season, data were aggregated into weeks (records taken within each Monday to Sunday period to account for differences in sampling day). These weekly data were also aggregated across subsamples, and subsample numbers were tallied to account for any variation in sampling effort due to lost traps. A typical sample represented all individuals of each target species captured each week within a treatment by repetition combination, with the total actual traps (usually 5) reported for this time recorded as a sampling effort covariate. When aggregated this way, typical captures were zero-biased, with a range of 0-70 (median=0, mean=1.46) beetles of a species per trapping unit, per week. Yearly data were compiled in a similar way, except data were aggregated across sampling weeks, and reported as total captures of a given species, per treatment by repetition combination within the year, with a covariate to account for sampling effort (~50 traps per year). This temporal resolution resulted in captures ranging from 0-198 individuals of a species per observation (median =8, mean =14.8). Data were culled at the first week of September (week of year = 35) as data collection usually ended by late August, making records beyond this point sparse in most years.

In addition to the species count data, we compiled contextual data from the weather station records available from the KBS site (https://lter.kbs.msu.edu/datatables/7). Because some missing measurements occurred in these data, we adapted a gap-filling algorithm which took the average of the proceeding and immediately following measurement, then substituted this value for any blank observations (Hermann et al. 2016). This manipulation allowed reasonable estimates of cumulative weather data metrics to be calculated.

Weather data were aggregated at complementary resolutions to accompany each temporal resolution of lady beetle data. For weekly data, we computed degree day and precipitation accumulation (after Hermann et al 2016) within each week, and then aggregated over the growing season, using daily maximum and minimum temperatures as inputs, baseline development threshold of 10°C, and a starting date of January 1. We also computed several derived weather metrics appropriate to the resolution of the insect data (number of rainy days within an observation week, minimum and maximum temperatures observed within that sampling week). For yearly data, we computed several derived metrics to characterize weather at a coarser level within a given year, through the growing season. For this, we used four time points (week of year 20, 25, 30, 35) and computed the degree day and precipitation accumulation at each, allowing us to examine how periods of weather of a given type affected the overall number of each species observed in a given year.

To examine patterns of habitat use across the nine plant communities between the two species by time period, we conducted 2-D non-metric multidimensional scaling (NMDS) analyses on each data resolution using *vegan v. 2.5-7* (Oksanen et al. 2013). First, data were subjected to square root transformation and Wisconsin double standardization, and then were subjected to a permutational ANOVA to determine if statistical differences occurred between the two species in their habitat use at either time period. Environmental variables were then fitted to the NMDS for each temporal resolution (Oksanen et al. 2013). Parameters used in the final model were selected from a global model that included all computed weather parameters, subject to backwards selection to simplify a final model with parameters with the strongest explanatory results.

To examine the within-season dynamics between the two lady beetle species and the roles of multiple environmental factors in driving abundances of each species, we constructed a generalized additive model for the number of lady beetles observed, where each parameter tested was allowed to interact with species, allowing us to directly examine how each species response to a given parameter differed. All model structures tested included an offset for trapping effort, used a quasi-Poisson error structure, and smoothing parameters were fitted using restricted maximum likelihood (REML) using *mgcv v. 1.8-36* (Wood 2006, Marra and Wood 2011). Because of strong autocorrelation between many weather parameters (Appendix S1: Figure S1), we used a substitution-based forward model selection approach by substituting a single parameter of each type (temperature variables: mean temperature, maximum temperature, minimum temperature, degree day accumulation within sampling week, total degree day accumulation; precipitation variables: mean daily precipitation, number of rainy days, precipitation accumulation over the year, maximum daily precipitation within sample week) and evaluating model performance for single variables, and then using the best single predictor model (as determined by lowest value of -REML). We then added terms with Pearson correlation <0.5 with existing terms to our best single-environmental variable model. Our final model included parameters for degree day accumulation, maximum daily rainfall within the sampling week, maximum temperature observed in the sampling week, and year, as well as a categorical variable for habitat of capture.

Between-year dynamics of the two species were modeled similarly to within-year dynamics, but to allow different conditions to act as drivers for each species, we constructed separate models and corresponding model selections for each species. All models contained offset terms to account for sampling effort, a categorical variable for habitat of capture, and used a quasi-Poisson error structure. Smoothing was performed using REML. Each of these models for the number of adults of a given species also included a covariate for the number of adults of the other species. Because an initial correlation analysis suggested minimal autocorrelation between environmental parameters at this scale (Pearson correlations of all parameters <0.5, Appendix S1: Figure S2) we completed a backward selection of all environmental variables, assessing model fit and concurvity at each step, eliminating the parameter with the least apparent contribution to explanatory power of the model, and rerunning and repeating assessments. This procedure was repeated until concurvity estimates for all parameters were <0.8 and there were no more parameters that could be removed without losing explanatory power of the model. For all environmental parameters tested, the smoothing parameter was set to sp=1 to minimize the impact of outlying data points on the overall trends observed and to prevent overfitting (Hunsicker et al., 2016).

All data manipulation and aggregation, as well as all exploratory and analytical approaches were completed in R 4.1.1 (R Development Core Team 2017). The analysis code and development history are available at https://github.com/ReproducibleQM/space_invader.

## Results

For conciseness within the results, we have abbreviated *Coccinella septempunctata* as C7 and *Harmonia axyridis* as HA. After culling data in the first week of September in all years and all data prior to 1994, after HA arrived, we observed nearly equivalent total captures of the two species, with 19,637 individuals of C7 and 20,412 of HA. Overall, mean abundance (±SD) per observation at the weekly resolution was 1.4± 3.8 for C7 and 1.5 ± 2.9 for HA, and 14.5±22.3 and 15.1±17.1 for C7 and HA at the yearly resolution respectively. Raw abundance and habitat use patterns varied by species but highly overlapped (Figure 1). At the initiation of the study, C7 dominated captures but HA increased until the mid-2000s, and then the two species appeared to undergo a cyclical switching off in dominance for the most recent 15 years of the study (Figure 1A). Both species were present in all habitats examined, although C7 was more commonly captured in wheat than HA, and HA was more common than C7 in all forest treatments (Figure 1B).

**Figure 1:**
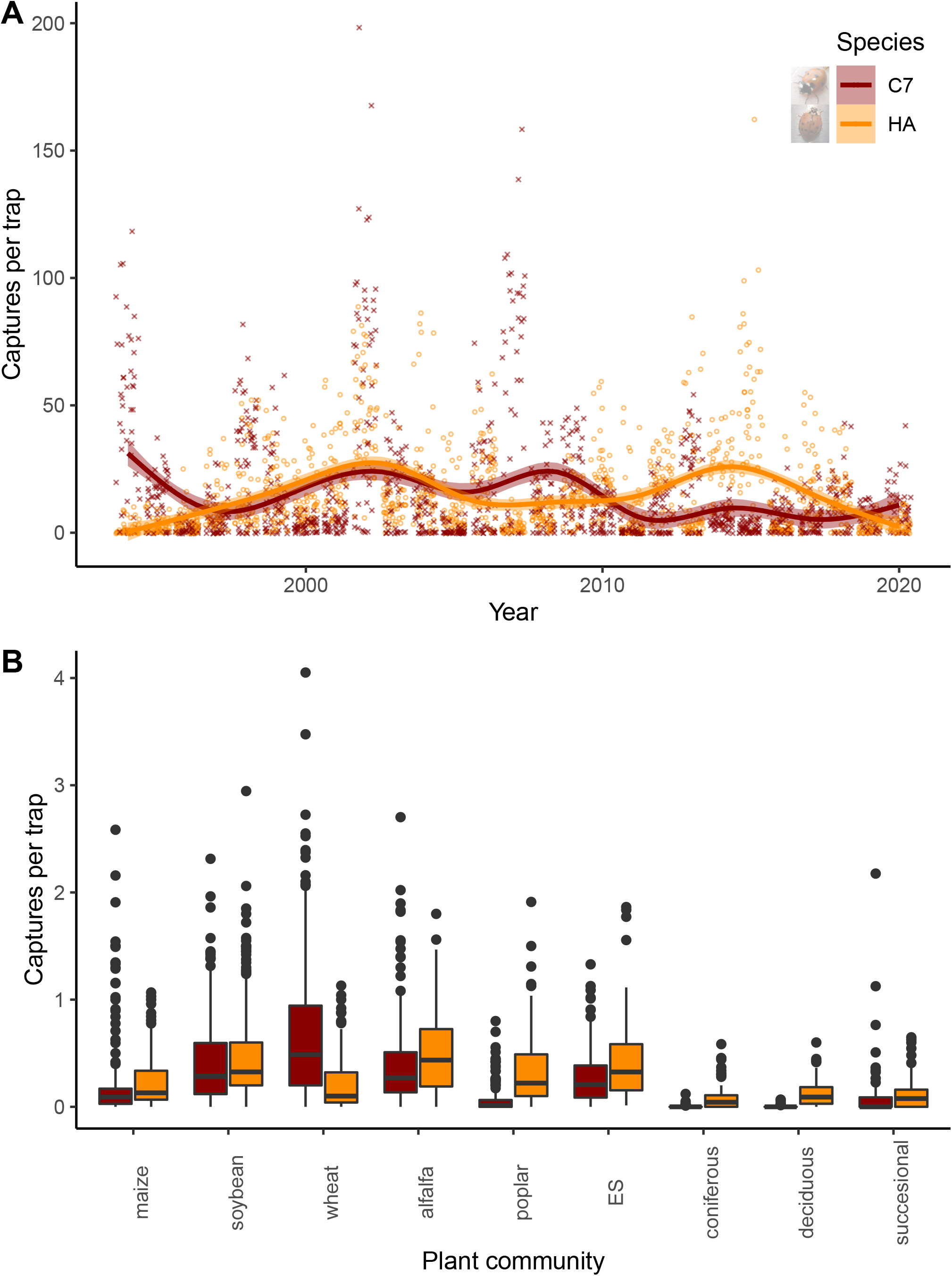
Raw trends in the abundance of two adventive lady beetle species at Kellogg Biological Station in Southwestern Michigan, 1994-2020. A) abundance of adult lady beetles, captured per repetition treatment combination (*ca*. 5 traps per week) B) the same data given by plant community treatment, aggregated over time. *Coccinella septempunctata* (C7) captures are given in red, *Harmonia axyridis* (HA) in orange. ES refers to the early successional vegetation treatment.

### Multivariate analysis

Habitat use patterns varied between the two species at both the yearly (F_1,322_=14.25, R^2^=0.04, *P*=0.001) and weekly (F_1,630_=21.93, R^2^=0.03, *P*=0.001) resolution. Non-metric multidimensional scaling (Figure 2) suggested a distinct clustering of spatio-temporal distribution between the species at both resolutions. Measures of precipitation and degree day accumulation were included in the best model for environmental drivers of the distribution of observations in both cases, suggesting these drivers both impacted within-season and between-season dynamics. At the yearly resolution, the final variables included in the environmental model were degree day accumulation at 35 weeks (r^2^=0.086, *P*=0.001), precipitation accumulation at 35 weeks (r^2^=0.064, *P*=0.001), and year (treated as a continuous variable; r^2^=0.025, *P*=0.017). At the weekly resolution, the environmental model included within-season degree day accumulation (r^2^=0.0328, *P*=0.001), within-season precipitation accumulation (r^2^=0.014, *P*=0.001), and year (r^2^=0.011, *P*=0.045). In both cases, year and precipitation accumulation plotted as nearly perfectly collinear, but moving in opposite directions. The degree day accumulation factor plotted with a larger orthogonal component, more closely corresponding to the axis of differentiation between the distributions of the two species, suggesting that while precipitation patterns are clearly changing at our study site, it is likely that temperature cues contribute more strongly to niche differentiation of these two species.

**Figure 2:**
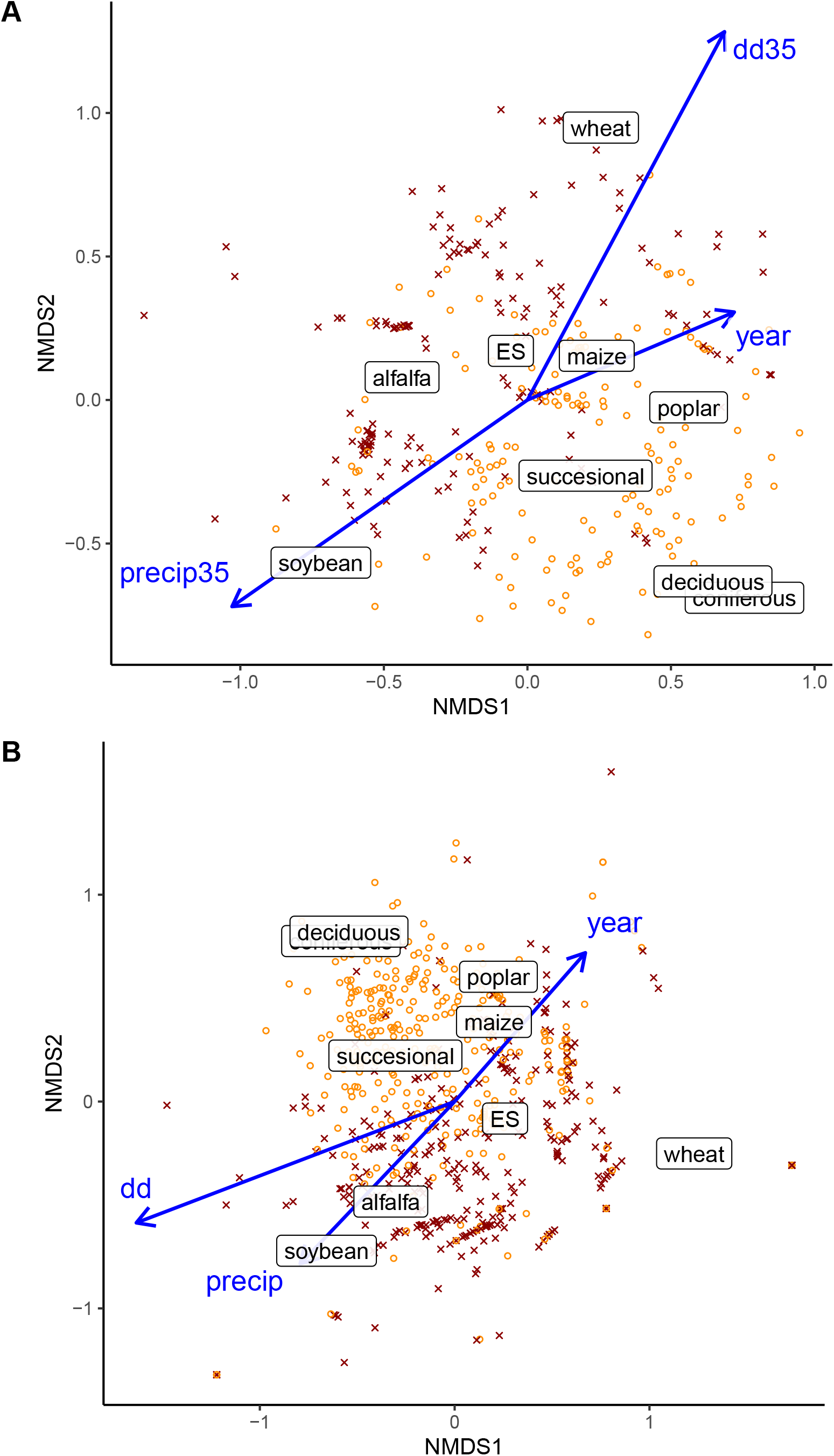
Non-metric multidimensional scaling of habitat use patterns by two adventive lady beetle species at Kellogg Biological Station in Southwestern Michigan, 1994-2020, at two temporal aggregations. A) yearly resolution, 2-D stress= 0.211; B) weekly resolution, 2-D stress= 0.226. *Coccinella septempunctata* observations are given by red X, *Harmonia axyridis* observations are given by orange O. Centroids for the plant community sites where insects were captured are plotted on both ordinations. ES refers to the early successional vegetation treatment. Vectors for environmental variables explaining significant variation in the community are labeled on the plot: dd refers to degree day accumulation over the year, dd35 is degree day accumulation at 35 weeks of the year; similarly ‘precip’ is the precipitation accumulation over the year, and ‘precip 35’ is the accumulation at 35 weeks.

### Within-season GAM model

The final model for within-season captures of lady beetles (-REML=26835, n=27432, Deviance explained = 36.4%) included terms for the categorical variables of habitat, species, and the interaction between habitat and species as well as smoothing terms for degree day accumulation, maximum rainfall within the sampling week, maximum temperature within the sampling week, and year, each interacting with species (Figure 3; Appendix S1: Table S1, S2). The majority of within-season variation in both species was attributable to habitat of capture and degree day accumulation: the habitat and year model explained 25% of deviance in the data, the model with degree day accumulation added to these terms accounted for 33.1% of the deviation, and habitat alone without any covariates accounted for 14.9%. Both species exhibited strong, differential within-season responses to degree day accumulation (Figure 3A). Using the instantaneous rate of change of the line of best fit of our model parameterized using ‘average’ environmental conditions in a single habitat (alfalfa), we predicted the approximate degree day accumulations where activity peaks occurred and magnitude of these peaks. The model predicted C7 captures ranging from 0 to 1.76 beetles per sample, with a mean capture rate of 0.77 beetles per sample. C7 had a large activity peak early in the season at approximately 550 degree days accumulated at an estimated 1.76 beetles per sample, and a slight peak later in the season at an accumulation of 1210 degree days and approximately 0.41 beetles per sample. The model predicted a range of 0 to 1.60 beetles per sample, with a mean of 0.85. HA had a major late season activity peak near 1250 degree days accumulation, with an estimated 1.55 beetles per sample, and populations remained high and growing thereafter, and we observed two lesser early season activity peaks near 410 and 695 degree days, with predicted captures of 0.81 and 0.67 beetles per sample, respectively. Although model selection favored the inclusion of terms describing the maximum rainfall and temperature the week when observations were taken, these terms explained relatively little variation: their addition increased the total deviance explained by the model to 36.4%. In both cases, the inclusion of these terms explained very little new variation for HA and suggested perhaps a slight increase in C7 activity in weeks that were moderately rainy (Figure 3B) or had warmer temperatures once the effect of degree day accumulation was accounted for (Figure 3C). Once other factors were controlled for in the model, residual habitat use patterns between the two species became strikingly differential (Figure 3D): C7 was much more variable in habitat use patterns. For example, in the perennial cropland habitats (alfalfa, early successional vegetation and poplar plantations), the model predicted 1.76 captures of C7 per sample in alfalfa, 1.31 captures in early successional vegetation to essentially no capture of this species in poplar plantations at its activity peak. Under these same conditions, the model predicted 0.65, 0.49 and 0.24 captures of HA in alfalfa, early successional vegetation, and poplar plantations respectively. For these same habitats at the HA activity peak, the model predicted 0.38 captures of C7 in alfalfa and essentially none in the other habitats, while HA captures were predicted to be 1.54, 1.38 and 1.13 for each habitat, respectively. Finally, similarly, year-to-year dynamics of both species still contributed considerable variation to the captures (Figure 3E).

**Figure 3:**
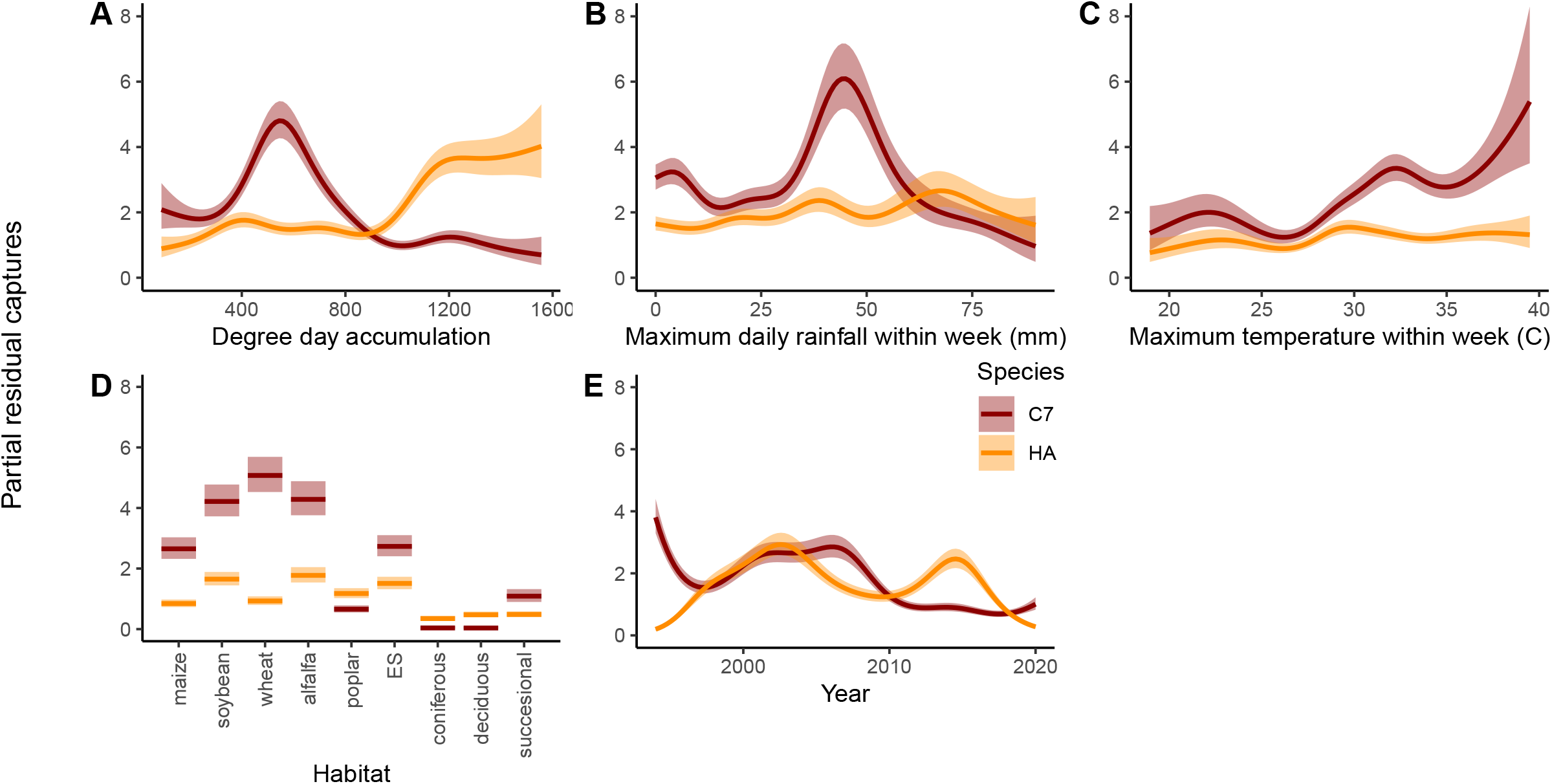
Partial effects of terms included in a within-season Generalized Additive Model for two adventive lady beetle species at Kellogg Biological Station in Southwestern Michigan, 1994-2020, using weekly observations of lady beetles captured in nine plant communities. A) Within-season degree day accumulation (base 10°C); B) Maximum rainfall observed within sampling week (mm)); C) maximum temperature observed during sampling week (degrees Celsius); D) residual habitat effects; and E) remaining year-to-year population variation. *Coccinella septempunctata* (C7) effects are given in red, *Harmonia axyridis* (HA) in orange.

### Between-year model

In the examination of between-year dynamics of the two species, model selection favored GAM models with three environmental parameters for both species (Appendix S1: Table S3), with none of these factors overlapping between models (Appendix S1: Table S4). The C7 model explained more variation in the year-to-year abundance of this species (-REML=1905, n=1353, Deviance explained = 79.0%) than the model for HA abundance (-REML=1855, n=1353, Deviance explained = 67.3%). In general, relationships between HA and environmental parameters were slight, and HA abundance peaked near the mean values of each of the three environmental parameters included in the model (Figure 4), suggesting that while HA is not affected strongly by environmental conditions, it generally is most abundant in years when conditions are near the mean. C7 was observed to have both stronger responses to environmental conditions, but also, was more negatively associated with mean conditions, i.e. having higher numbers in years associated with cool early summer conditions and warmer, drier conditions late in the growing season. A spurious peak was observed for very low degree day accumulations at 35 weeks in the C7 model (Figure 4): this peak was driven by a single observation at the extreme of the distribution for this parameter (Appendix S1: Figure S2).

**Figure 4:**
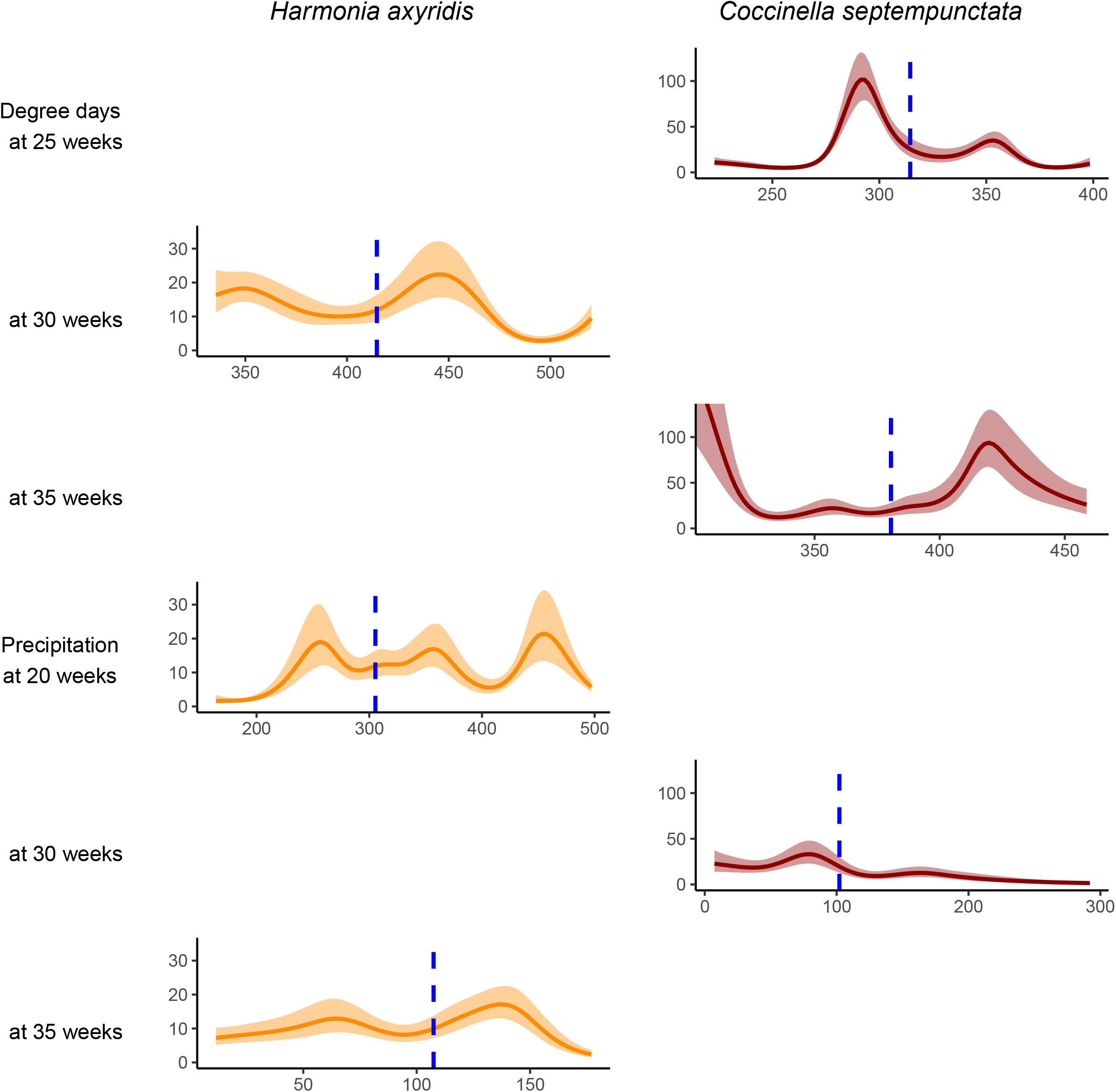
Partial effects of environmental parameters from a between-season Generalized Additive Models for two adventive lady beetle species at Kellogg Biological Station in Southwestern Michigan, 1994-2020. Each species was modeled and subjected to model selection separately. *Coccinella septempunctata* (C7) model effects are given in red, *Harmonia axyridis* (HA) in orange. Models included term for between-habitat variation.

## Discussion

Following their establishment and spread within North America, *H. axyridis* and *C. septempunctata* have been commonly collected within similar environments, but how these species are able to coexist is poorly understood. In this study, we used a 31-year dataset from an agricultural landscape in southwestern Michigan, USA to understand mechanisms of coexistence among these dominant species. We found evidence of long-term coexistence, as both species were collected in similar abundance overall and found in all habitat types. Our findings indicated that a combination of small-scale niche partitioning via environmental, habitat, and seasonal mechanisms contributed to coexistence among these species. While net populations of species observed over the course of the study were nearly identical, populations of *C. septempunctata* were more variable. Within the midwestern US, *H. axyridis* seems to have broader habitat and abiotic environmental preferences, while *C. septempunctata* tends to thrive under more specific ecological conditions which are not the average for this region.

Following the arrival and establishment of *H. axyridis* within the region, these two species experienced patterns of cyclical dominance in abundance across years, suggesting there is some degree of overlap in their niches. This finding aligns with research documenting overlap in preferences for aphid prey among the two species (Koch 2003, Hodek and Michaud 2008, Roy et al. 2016) as well as evidence of antagonistic competitive interactions in some environments (Snyder 2009, Gagnon et al. 2011). These patterns of between year partitioning may suggest coexistence via the storage effect. The storage effect hypothesis posits that competitors may coexist if overlapping generations experience temporal fluctuations in the recruitment of individuals due to species-specific responses to the environment (Chesson and Huntly 1997, Chesson 2000b). Yearly fluctuations in environmental conditions may have differentially benefited one species over the other, resulting in patterns of alternating dominance. Environmental fluctuations may have directly impacted these lady beetles through differential species’ preferences or indirectly via changes in prey populations.

Although we were unable to distinguish between direct and indirect mechanisms, populations of *H. axyridis* and *C. septempunctata* responded differentially to precipitation and temperature. Abundance of *C. septempunctata* was negatively associated with means of several environmental metrics, as populations were more successful in years characterized by cooler temperatures early in the growing season as well as warmer temperatures and drier conditions in late summer. In contrast, *H. axyridis* was not strongly affected and was generally most abundant in years when temperatures and the amount of precipitation were near the mean. *Harmonia axyridis* has a broad global distribution and can be found at a range of altitudes, but is generally more abundant in cool, mesic climates and less common in the tropics (Roy et al. 2016). However, this species typically overwinters in aggregations in hibernacula such as buildings and under tree bark, which may buffer the effects of cold temperatures to some extent (Roy et al. 2016). In contrast, *C. septempunctata* has been observed overwintering in the soil and leaf litter layers of forests and tree margins (Turnock and Wise 2004), suggesting that cool-weather tolerance in this species may contribute to differential success within a given year. Cooler conditions in early summer may favor *C. septempunctata* via temperature dependence in foraging rates. While both species have similar prey attack rates at 26°C (Xue et al. 2009), attack rates on aphids were at their highest at lower temperatures (around 20°C) for *C. septempunctata*(Khan and Khan, 2010). Meanwhile, predation rates by *H. axyridis* increased with increasing temperature up to 35°C, and were markedly decreased below 25°C (Islam et al 2021). Both species had positive associations with warmer-than-average temperatures in mid or late summer, which could be associated with temperature-modified activity of prey aphids (Crossley et al. 2022). Within this region of the midwestern US, direct and/or indirect responses of *H. axyridis* and *C. septempunctata* to temperature and precipitation have contributed to niche differences and cyclical patterns of dominance over time.

Within an average year, the population dynamics of *H. axyridis* and *C. septempunctata* showed differing responses to seasonal change, as abundances of each species peaked at different times during the growing season. Both species responded strongly to degree day accumulation, with the abundance of *C. septempunctata* peaking earlier, while *H. axyridis* became more abundant later in the growing season. This finding contradicts predictions that high overwintering survival of *H. axyridis* will lead to larger populations in the spring than *C. septempunctata* (Raak-van der Berg et al. 2012). The differing phenological responses likely allow these two competitors to locally coexist on short time scales by creating temporal niche differences. Differences in phenology influence when and at which developmental stage species interactions occur, and thus patterns of coexistence (Rudolf 2019, Blackford et al. 2020). Both *H. axyridis* and *C. septempunctata* require aphid prey to some degree to successfully develop and reproduce (Berkvens et al. 2008, Hodek and Michaud 2008; Zaviezo et al. 2019) such that their phenology may be strongly tied to thresholds in prey resources. For example, at low aphid densities, reproduction of *H. axyridis* did not occur or was very low in the laboratory than at high aphid densities (Zaviezo et al. 2019). In Europe where *C. septempunctata* is native, this species exploits immigrating aphid populations in cereals beginning in May until the crop matures in July (Honek et al. 2019), suggesting a tight association with this prey resource. In contrast, the breeding period of *H. axyridis* is greatly extended, and populations will continue to feed and grow into late summer risking incomplete development of later generations (Honek et al. 2018). Responses of *H. axyridis* to populations of soybean aphid suggest that late season abundance was linked to prey availability early in the growing season (Bahlai et al. 2015).

Although phenological differences among species may promote local coexistence through increased niche differences (i.e. stabilizing mechanisms) (Albrecht and Gotelli 2001), it is also possible that greater phenological differences can lead to increased fitness differences (i.e. equalizing mechanisms) that promote competitive exclusion (Godoy and Levine 2014, Blackford et al. 2020). For instance, in years where *C. septempunctata* activity is favored early in the growing season, this species may deplete local prey resources (Bianchi and Van der Werf 2004), negatively impacting early generations of *H. axyridis.*

Patterns of abundance of *H. axyridis* and *C. septempunctata* varied among plant communities, indicating spatial niche partitioning among habitat types. Abundance of *C. septempunctata* was highest in soybean, wheat, and alfalfa, which suggests a habitat preference for these crop types. Other studies have reported apparent associations between *C. septempunctata* and various herbaceous crop environments, including cereals (Honek et al. 2014, 2019), potato (Alyokhin and Sewell 2004), alfalfa, and maize (Elliott et al. 1996). Abundance of *H. axyridis* was more consistent across all plant communities, which aligns with studies that have identified this exotic species as having a broad habitat range that includes agricultural fields, orchards, vineyards, parks, and residential yards and gardens (Koch 2003, Adriaens et al. 2008; Roy et al. 2016). Although *H. axyridis* did not show a preference for a particular habitat type, this species was more abundant than *C. septempunctata* in forests. Populations of *H. axyridis* have been commonly reported in arboreal habitats in Europe (Adriaens et al. 2008, Vandereycken et al. 2012, Honek et al. 2014, Panigaj et al. 2014), wherein their abundance was found to be 7.5x higher on trees than other herbaceous plants including crops (Honek et al. 2019). Larvae of *H. axyridis* are morphologically adapted for exploiting canopy environments compared to *C. septempunctata* due to their well-developed anal disc for adhering to plant surfaces (Eigenbrode et al. 2009).

Although *C. septempunctata* displayed clear preferences for crops, *H. axyridis* also was collected in these habitats. There is some evidence that when these two species overlap in habitat, they partition resources at finer spatial scales. For example, *C. septempunctata* did not change its habitat use patterns on apple trees in the presence of *H. axyridis,* and instead, these species limited niche overlap by vertically partitioning resources (Lucas et al. 2002). Moreover, *C. septempunctata* has been shown to remain in preferred habitats such as alfalfa and soybean for extended periods, while *H. axyridis* frequently disperses among habitats especially later in the growing season (Forbes and Gratton 2011). Lady beetle activity and thus, the interactions between these two dominant species were strongly related to temperatures via daily degree day accumulation, and to a lesser extent, precipitation accumulation, suggesting climate change may alter these coexistence mechanisms. Phenological responses of species and the seasonal timing of interactions are not static but vary yearly with environmental conditions (Singer and Parmesan 2010, Rudolf 2019). Climate change is causing gradual increases in temperature on a global scale as well as altered patterns of precipitation (IPCC 2014). Warmer temperatures affect lady beetles directly by increasing metabolism and rates of egg and larval development, but also require higher rates of prey consumption (Banfield-Zanin and Leather 2016, Speights and Barton 2019). Above a heat stress threshold, increased temperatures may cause high mortality of eggs, larvae, and some adults (Acar et al. 2001, Knapp and Nedvěd 2013). Prey responses to climate change also are complex and can be affected by physiological tolerances and host plant quality, which may indirectly affect interactions with lady beetles (Honek et al. 2017, Sloggett 2021). Responses of aphids are often species-specific but warming temperatures and milder winters generally shift aphid phenology such that early season flight activity occurs sooner in the season (Bell et al. 2015, Wu et al. 2020). Under laboratory simulated drought conditions, aphids tended to be less nutritious and smaller in size, requiring higher consumption rates by lady beetles to meet dietary requirements (Banfield-Zanin and Leather 2016). Therefore, climate change may lead to directional shifts in phenology and timing of trophic interactions such that synchrony of predator-prey interactions may be disrupted, with implications for biological control. Adding complexity, continued promotion of landscape simplification rather than diversification may interact with climate change to influence coexistence outcomes among lady beetle species (Schulte et al. 2021). Opportunistic generalist species such as *H. axyridis* may be well adapted to respond to environmental shifts, providing a competitive advantage.

Coexistence of these dominant exotic lady beetle species may be further affected by other mechanisms that were not investigated in this study such as differential predation and parasitism. For example, the enemy release hypothesis posits that exotic species will experience reduced top-down control from natural enemies in their introduced range, leading to successful establishment and spread (Keane and Crawley 2002, Shea and Chesson 2002, Colautti et al. 2004; Roy et al. 2011). For instance, there is evidence that *H. axyridis* is less susceptible to the hymenopteran parasitoid *Dinocampus coccinellae* (Schrank) that commonly parasitizes *C. septempunctata* in Europe (Geoghegan et al. 1998, Berkvens 2010, Comont et al. 2014).These effects may manifest at the population level and furthermore, would likely interact with habitat and environmental partitioning observed in this study.

In this study, we use trap captures as a proximal measurement to represent the dynamics of the species under examination, these traps more meaningfully capture activity density of lady beetles: insects moving through the environment that are drawn to, or collide with traps. It is important to note that, while this trapping method is advantageous in this context because it is inexpensive and relatively easy to deploy in a consistent way over years of sampling, sticky cards are also prone to inherent biases. For instance, yellow sticky cards may have differential attractiveness to different species of ladybeetle (Musser et al. 2004), and like many trapping methods, may have differing capture efficiency depending on the habitats in which they are deployed (Missa et al. 2009).

Successful establishment and spread of exotic species are influenced by a variety of factors including local niche processes and interactions among species within the native community. Although research has largely examined biological invasions in the context of their impacts on native species, communities are frequently invaded by multiple exotic species which then interact with each other to influence establishment success. Exotic lady beetles *H. axyridis* and *C. septempunctata* were introduced into North America for biological control in agricultural systems (Obrycki and Kring 1998, Evans 2009, Rondoni et al. 2021), and these species have since become dominant within many native communities. Research has documented asymmetric competitive interactions in favor of *H. axyridis,* which suggests other forms of niche partitioning contribute to coexistence among these dominant invaders. Using a 31-year dataset from southwestern Michigan, USA, we documented evidence of long-term coexistence of these exotic lady beetle species via temporal and spatial niche partitioning occurring within and across years. Our findings indicated that within this region *H. axyridis* has broader habitat and abiotic environmental preferences, while *C. septempunctata* thrives under more specific ecological conditions. Ecological differences among these exotic species have promoted coexistence through environmental, seasonal, and habitat niche partitioning within agricultural landscapes in the midwestern US. Understanding mechanisms that allow coexistence of dominant exotic species contributes to native biodiversity conservation in the face of global change.

## Acknowledgements

This manuscript was prepared by students in CB’s Reproducible Quantitative Methods graduate class at Kent State University. The Mozilla Foundation supported CB during the development of this course. We thank Lama Tawk for leading a discussion of coexistence theory in the class. Support for this research was provided by National Science Foundation grants OAC 1838807 and DBI 2045721 to CB, NSF Long-term Ecological Research Program (DEB 1832042) at the Kellogg Biological Station, and Michigan State University AgBioResearch. DAL acknowledges support from The Great Lakes Bioenergy Research Center, U.S. Department of Energy, Office of Science, Office of Biological and Environmental Research (Award DE-SC0018409). The long-term lady beetle experiment at KBS-LTER was initiated in 1989 by Stuart Gage and Manuel Colunga-Garcia. We sincerely thank the ongoing efforts of the research staff supporting the long-term study, particularly: Stacey Van Der Wulp, Sven Bohm, Julia Perrone, Elizabeth D’Auria, and countless undergraduate assistants over the project’s history. Two reviewers provided helpful suggestions that improved the manuscript.

## Literature Cited

Acar, E. B., B. N. Smith, L. D. Hansen, and G. M. Booth. 2001. Use of calorespirometry to determine effects of temperature on metabolic efficiency of an insect. Environmental Entomology. 30: 811–816.

Adriaens, T., G. San Martin y Gomez, and D. Maes. 2008. Invasion history, habitat preferences and phenology of the invasive ladybird *Harmonia axyridis* in Belgium, pp. 69–88. *In* Roy, H.E., Wajnberg, E. (eds.), From Biological Control to Invasion: The Ladybird Harmonia axyridis as a Model Species. Springer Netherlands, Dordrecht.

Albrecht, M., and N. J. Gotelli. 2001. Spatial and temporal niche partitioning in grassland ants. Oecologia. 126: 134–141.

Alyokhin, A., and G. Sewell. 2004. Changes in a lady beetle community following the establishment of three alien species. Biological Invasions. 6: 463–471.

Bahlai, C. A., M. Colunga-Garcia, S. H. Gage, and D. A. Landis. 2013. Long term functional dynamics of an aphidophagous coccinellid community are unchanged in response to repeated invasion. PLoS One. 8: e83407.

Bahlai, C. A., M. Colunga-Garcia, S. H. Gage, and D. A. Landis. 2015. The role of exotic ladybeetles in the decline of native ladybeetle populations: evidence from long-term monitoring. Biological Invasions. 17: 1005–1024.

Bahlai, C. A., C. Hart, M. T. Kavanaugh, J. D. White, R. W. Ruess, T. J. Brinkman, H. W. Ducklow, D. R. Foster, W. R. Fraser, H. Genet, P. M. Groffman, S. K. Hamilton, J. F. Johnstone, K. Kielland, D. A. Landis, M. C. Mack, O. Sarnelle, and J. R. Thompson. 2021. Cascading effects: insights from the U.S. Long Term Ecological Research Network. Ecosphere. 12: e03430.

Bahlai, C. A., W. vander Werf, M. O’Neal, L. Hemerik, and D. A. Landis. 2015. Shifts in dynamic regime of an invasive lady beetle are linked to the invasion and insecticidal management of its prey. Ecological Applications. 25: 1807–1818.

Banfield-Zanin, J. A., and S. R. Leather. 2016. Prey-mediated effects of drought on the consumption rates of coccinellid predators of *Elatobium abietinum*. Insects. 7.

Bell, J. R., L. Alderson, D. Izera, T. Kruger, S. Parker, J. Pickup, C. R. Shortall, M. S. Taylor, P. Verrier, and R. Harrington. 2015. Long-term phenological trends, species accumulation rates, aphid traits and climate: five decades of change in migrating aphids. Journal of Animal Ecology. 84: 21–34.

Berkvens, N., J. Bonte, D. Berkvens, K. Deforce, L. Tirry, and P. De Clercq. 2008. Pollen as an alternative food for *Harmonia axyridis*, pp. 201–210. *In* Roy, H.E., Wajnberg, E. (eds.), From Biological Control to Invasion: The Ladybird Harmonia axyridis as a Model Species. Springer Netherlands, Dordrecht.

Berkvens, N., C. Landuyt, K. Deforce, D. Berkvens, L. Tirry, and P. De Clercq. 2010. Alternative foods for the multicoloured Asian lady beetle *Harmonia axyridis* (Coleoptera: Coccinellidae). Eur. J. Entomol. 107: 189–195.

Berkvens, N., J. Moens, D. Berkvens, M. A. Samih, L. Tirry, and P. De Clercq. 2010. *Dinocampus coccinellae* as a parasitoid of the invasive ladybird *Harmonia axyridis* in Europe. Biological control. 53: 92–99.

Bianchi, F. J. J. A., and W. Van der Werf. 2004. Model evaluation of the function of prey in non-crop habitats for biological control by ladybeetles in agricultural landscapes. Ecological Modelling. 171: 177–193.

Blackburn, T. M., P. Pyšek, S. Bacher, J. T. Carlton, R. P. Duncan, V. Jarošík, J. R. U. Wilson, and D. M. Richardson. 2011. A proposed unified framework for biological invasions. Trends in Ecology & Evolution. 26: 333–339.

Blackford, C., R. M. Germain, and B. Gilbert. 2020. Species differences in phenology shape coexistence. The American Naturalist. 195: E168–E180.

Boettner, G. H., J. S. Elkinton, and C. J. Boettner. 2000. Effects of a biological control introduction on three nontarget native species of saturniid moths. Conservation Biology. 14: 1798–1806.

Bradshaw, C. J. A., B. Leroy, C. Bellard, D. Roiz, C. Albert, A. Fournier, M. Barbet-Massin, J.-M. Salles, F. Simard, and F. Courchamp. 2016. Massive yet grossly underestimated global costs of invasive insects. Nature Communications. 7: 12986.

Cheng, J., P. Li, Y. Zhang, Y. Zhan, and Y. Liu. 2020. Quantitative assessment of the contribution of environmental factors to divergent population trends in two lady beetles. Biological Control. 145: 104259.

Chesson, P. 2000a. Mechanisms of maintenance of species diversity. Annual Review of Ecology and Systematics. 31: 343–366.

Chesson, P. 2000b. General theory of competitive coexistence in spatially-varying environments. Theoretical Population Biology. 58: 211–237.

Chesson, P., and N. Huntly. 1997. The roles of harsh and fluctuating conditions in the dynamics of ecological communities. The American Naturalist. 150: 519–553.

Colautti, R. I., A. Ricciardi, I. A. Grigorovich, and H. J. MacIsaac. 2004. Is invasion success explained by the enemy release hypothesis? Ecology letters. 7: 721–733.

Colunga-Garcia, M., and S. H. Gage. 1998. Arrival, establishment, and habitat use of the multicolored Asian lady beetle (Coleoptera: Coccinellidae) in a Michigan landscape. Environmental Entomology. 27: 1574–1580.

Colunga-Garcia, M., S. H. Gage, and D. A. Landis. 1997. Response of an assemblage of Coccinellidae (Coleoptera) to a diverse agricultural landscape. Environmental Entomology. 26: 797–804.

Comont, R. F., B. V. Purse, W. Phillips, W. E. Kunin, M. Hanson, O. T. Lewis, R. Harrington, C. R. Shortall, G. Rondoni, and H. E. Roy. 2014. Escape from parasitism by the invasive alien ladybird, *Harmonia axyridis*. Insect Conservation and Diversity. 7: 334–342.

Crossley, M.S., Lagos Kutz, D., Davis, T.S., Eigenbrode, S.D., Hartman, G.L., Voegtlin, D.J. and Snyder, W.E., 2022. Precipitation change accentuates or reverses temperature effects on aphid dispersal. Ecological Applications, p.e2593.

Eigenbrode, S. D., W. E. Snyder, G. Clevenger, H. Ding, and S. N. Gorb. 2009. Variable attachment to plant surface waxes by predatory insects, pp. 157–181. *In* Gorb, S.N. (ed.), Functional Surfaces in Biology: Adhesion Related Phenomena Volume 2. Springer Netherlands, Dordrecht.

Elliott, N., R. Kieckhefer, and W. Kauffman. 1996. Effects of an invading coccinellid on native coccinellids in an agricultural landscape. Oecologia. 105: 537–544.

Evans, E. W. 2004. Habitat displacement of North American ladybirds by an introduced species. Ecology. 85: 637–647.

Evans, E. W. 2009. Lady beetles as predators of insects other than Hemiptera. Biological Control. 51: 255–267.

Forbes, K. J., and C. Gratton. 2011. Stable isotopes reveal different patterns of inter-crop dispersal in two ladybeetle species. Ecological Entomology. 36: 396–400.

Gagnon, A.-È., G. E. Heimpel, and J. Brodeur. 2011. The ubiquity of intraguild predation among predatory arthropods. PLOS ONE. 6: e28061.

Gandhi, K. J. K., and D. A. Herms. 2010. Direct and indirect effects of alien insect herbivores on ecological processes and interactions in forests of eastern North America. Biological Invasions. 12: 389–405.

Gardiner, M. M., K. I. Perry, C. B. Riley, K. J. Turo, Y. A. Delgado de la flor, and F. S. Sivakoff. 2021. Community science data suggests that urbanization and forest habitat loss threaten aphidophagous native lady beetles. Ecology and Evolution. 11: 2761–2774.

Geoghegan, I. E., T. M. O. Majerus, and M. E. Majerus. 1998. Differential parasitisation of adult and pre-imaginal *Coccinella septempunctata* (Coleoptera: Coccinellidae) by *Dinocampus coccinellae* (Hymenoptera: Braconidae). European Journal of Entomology. 95: 571–579.

Godoy, O. 2019. Coexistence theory as a tool to understand biological invasions in species interaction networks: Implications for the study of novel ecosystems. Functional Ecology. 33: 1190–1201.

Godoy, O., and J. M. Levine. 2014. Phenology effects on invasion success: insights from coupling field experiments to coexistence theory. Ecology. 95: 726–736.

Grez, A. A., T. A. Rand, T. Zaviezo, and F. Castillo-Serey. 2013. Land use intensification differentially benefits alien over native predators in agricultural landscape mosaics. Diversity and Distributions. 19: 749–759.

Harmon, J. P., E. Stephens, and J. Losey. 2007. The decline of native coccinellids (Coleoptera: Coccinellidae) in the United States and Canada, pp. 85–94. *In* New, T.R. (ed.), Beetle Conservation. Springer Netherlands, Dordrecht.

Hermann, S. L., S. Xue, L. Rowe, E. Davidson-Lowe, A. Myers, B. Eshchanov, and C. A. Bahlai. 2016. Thermally moderated firefly activity is delayed by precipitation extremes. Royal Society open science. 3: 160712.

HilleRisLambers, J., P. B. Adler, W. S. Harpole, J. M. Levine, and M. M. Mayfield. 2012. Rethinking community assembly through the lens of coexistence theory. Annu. Rev. Ecol. Evol. Syst. 43: 227–248.

Hironori, Y., and S. Katsuhiro. 1997. Cannibalism and interspecific predation in two predatory ladybirds in relation to prey abundance in the field. Entomophaga. 42: 153–163.

Hodek, I., and J. P. Michaud. 2008. Why is *Coccinella septempunctata* so successful? (A point-of-view). EJE. 105: 1–12.

Holway, D. A., L. Lach, A. V. Suarez, N. D. Tsutsui, and T. J. Case. 2002. The causes and consequences of ant invasions. Annu. Rev. Ecol. Syst. 33: 181–233.

Honek, A., A. F. Dixon, A. O. Soares, J. Skuhrovec, and Z. Martinkova. 2017. Spatial and temporal changes in the abundance and composition of ladybird (Coleoptera: Coccinellidae) communities. Current Opinion in Insect Science. 20: 61–67.

Honek, A., Z. Martinkova, A. F. G. Dixon, J. Skuhrovec, H. E. Roy, M. Brabec, and S. Pekar. 2018. Life cycle of *Harmonia axyridis* in central Europe. BioControl. 63: 241–252.

Honek, A., Z. Martinkova, P. Kindlmann, O. M. C. C. Ameixa, and A. F. G. Dixon. 2014. Long-term trends in the composition of aphidophagous coccinellid communities in Central Europe. Insect Conservation and Diversity. 7: 55–63.

Honek, A., Z. Martinkova, H. E. Roy, A. F. G. Dixon, J. Skuhrovec, S. Pekár, and M. Brabec. 2019. Differences in the phenology of *Harmonia axyridis* (Coleoptera: Coccinellidae) and native coccinellids in central Europe. Environmental Entomology. 48: 80–87.

Hunsicker, M.E., Kappel, C.V., Selkoe, K.A., Halpern, B.S., Scarborough, C., Mease, L., Amrhein, A., 2016. Characterizing driver–response relationships in marine pelagic ecosystems for improved ocean management. Ecological Applications 26, 651–663. https://doi.org/10.1890/14-2200

IPCC. 2014. Climate Change 2014: Synthesis Report. Contribution of Working Groups I, II and III to the Fifth Assessment Report of the Intergovernmental Panel on Climate Change. IPCC, Geneva, Switzerland.

Islam, Y., Shah, F.M., Rubing, X., Razaq, M., Yabo, M., Xihong, L. and Zhou, X., 2021. Functional response of *Harmonia axyridis* preying on *Acyrthosiphon pisum* nymphs: the effect of temperature. Scientific Reports, 11(1), pp.1–13.

Jeschke, J. M. 2014. General hypotheses in invasion ecology. Diversity and Distributions. 20: 1229–1234.

Keane, R. M. and M. J. Crawley. 2002. Exotic plant invasions and the enemy release hypothesis. Trends in Ecology & Evolution. 17: 164–170.

Kenis, M., M.-A. Auger-Rozenberg, A. Roques, L. Timms, C. Péré, M. J. W. Cock, J. Settele, S. Augustin, and C. Lopez-Vaamonde. 2009. Ecological effects of invasive alien insects. Biol Invasions. 11: 21–45.

Khan, M.R. and Khan, M.R., 2010. The relationship between temperature and the functional response of *Coccinella septempunctata* (L.)(Coleoptera: Coccinellidae). Pak. J. Zool, 42(4), pp.461–466.

Klooster, W. S., K. J. K. Gandhi, L. C. Long, K. I. Perry, K. B. Rice, and D. A. Herms. 2018. Ecological impacts of emerald ash borer in forests at the epicenter of the invasion in North America. Forests. 9.

Knapp, M., and O. Nedvěd. 2013. Gender and timing during ontogeny matter: Effects of a temporary high temperature on survival, body size and colouration in *Harmonia axyridis*. PLOS ONE. 8: e74984.

Koch, R. L. 2003. The multicolored Asian lady beetle, *Harmonia axyridis:* A review of its biology, uses in biological control, and non-target impacts. Journal of Insect Science. 3: 32.

Landis, D. 2020. Insect Population Dynamics on the Main Cropping System Experiment at the Kellogg Biological Station, Hickory Corners, MI (1989 to 2019) ver 30. Environmental Data Initiative. https://portal.edirepository.org/nis/mapbrowse?packageid=knb-lter-kbs.23.30

Leppanen, C., A. Alyokhin, and S. Gross. 2012. Competition for aphid prey between different lady beetle species in a laboratory arena. Psyche. 2012: 1–9. doi:10.1155/2012/890327

Levine, J. M., and C. M. D’Antonio. 1999. Elton revisited: A review of evidence linking diversity and invasibility. Oikos. 87: 15–26.

Liebhold, A. M., W. L. MacDonald, D. Bergdahl, and V. C. Mastro. 1995. Invasion by exotic forest pests: A threat to forest ecosystems. Forest Science. 41: a0001–z0001.

Lockwood, J. L., M. F. Hoopes, and M. P. Marchetti. 2013. Invasion Ecology. John Wiley & Sons.

Louda, S. M., D. Kendall, J. Connor, and D. Simberloff. 1997. Ecological effects of an insect introduced for the biological control of weeds. Science. 277: 1088–1090.

Lucas, É., I. Gagné, and D. Coderre. 2002. Impact of the arrival of *Harmonia axyridis* on adults of *Coccinella septempunctata* and *Coleomegilla maculata* (Coleoptera: Coccinellidae). Eur. J. Entomol. 99: 457–463.

Maredia, K., S. Gage, D. Landis, and T. Wirth. 1992. Ecological observations on predatory Coccinellidae (Coleoptera) in southwestern Michigan. The Great Lakes Entomologist. 25: 265–270.

Marra, G., and S.N. Wood. 2011. Practical variable selection for generalized additive models. Computational Statistics & Data Analysis. 55: 2372–2387.

Morin, R. S., A. M. Liebhold, P. C. Tobin, K. W. Gottschalk, and E. Luzader. 2007. Spread of beech bark disease in the eastern United States and its relationship to regional forest composition. Can. J. For. Res. 37: 726–736.

Missa, O., Y. Basset, A. Alonso, S. E. Miller, G. Curletti, M. De Meyer, C. Eardley, M. W. Mansell, and T. Wagner. 2009. Monitoring arthropods in a tropical landscape: relative effects of sampling methods and habitat types on trap catches. Journal of Insect conservation, 13: 103–118.

Musser, F. R., J. P. Nyrop, A. M. Shelton. 2004. Survey of predators and sampling method comparison in sweet corn. J. Econ. Entomol. 97: 136–144. https://doi.org/10.1093/jee/97.1.136

Obrycki, J. J., and T. J. Kring. 1998. Predaceous Coccinellidae in biological control. Annu. Rev. Entomol. 43: 295–321.

Oksanen, J., F. G. Blanchet, R. Kindt, M. J. Oksanen, and M. Suggests. 2013. Package ‘vegan.’ Community ecology package Version. 2: 0–0.

Panigaj, L., P. Zach, A. Honěk, O. Nedvěd, J. Kulfan, Z. Martinková, D. Selyemová, S. Viglášová, and H. E. Roy. 2014. The invasion history, distribution and colour pattern forms of the harlequin ladybird beetle *Harmonia axyridis* (Pall.) (Coleoptera, Coccinellidae) in Slovakia, Central Europe. Zookeys. 89–112.

Pell, J. K., J. Baverstock, H. E. Roy, R. L. Ware, and M. E. N. Majerus. 2008. Intraguild predation involving *Harmonia axyridis:* a review of current knowledge and future perspectives. BioControl. 53: 147–168.

Pyšek, P., P.E. Hulme, D. Simberloff, S. Bacher, T. M. Blackburn, J. T. Carlton, W. Dawson, F. Essl, L. C. Foxcroft, P. Genovesi, J. M. Jeschke, I. Kühn, A. M. Liebhold, N. E. Mandrak, L. A. Meyerson, A. Pauchard, J. Pergl, H. E. Roy, H. Seebens, H., M. van Kleunen, M. Vilà, M. J. Wingfield, and D. M. Richardson. 2020. Scientists’ warning on invasive alien species. Biological Reviews. 95: 1511–1534. https://doi.org/10.1111/brv.12627

R Development Core Team. 2017. R: A Language and Environment for Statistical Computing 3.3.3. R Foundation for Statistical Computing.

Raak-van den Berg, C. L., J. M. Stam, P. W. de Jong, L. Hemerik, and J. C. van Lenteren. 2012. Winter survival of *Harmonia axyridis* in The Netherlands. Biological Control, 60: 68–76.

Ricciardi, A. 2007. Are modern biological invasions an unprecedented form of global change? Conservation Biology. 21: 329–336.

Ricciardi, A., M. F. Hoopes, M. P. Marchetti, and J. L. Lockwood. 2013. Progress toward understanding the ecological impacts of nonnative species. Ecological Monographs. 83: 263–282.

Rondoni, G., I. Borges, J. Collatz, E. Conti, A. C. Costamagna, F. Dumont, E. W. Evans, A. A. Grez, A. G. Howe, E. Lucas, J.-É. Maisonhaute, A. Onofre Soares, T. Zaviezo, and M. J. W. Cock. 2021. Exotic ladybirds for biological control of herbivorous insects – a review. Entomologia Experimentalis et Applicata. 169: 6–27.

Roy, H. E., L. J. Lawson Handley, K. Schönrogge, R. L. Poland, and B. V. Purse. 2011. Can the enemy release hypothesis explain the success of invasive alien predators and parasitoids? BioControl. 56: 451–468.

Roy, H. E., P. M. J. Brown, T. Adriaens, N. Berkvens, I. Borges, S. Clusella-Trullas, R. F. Comont, P. De Clercq, R. Eschen, A. Estoup, E. W. Evans, B. Facon, M. M. Gardiner, A. Gil, A. A. Grez, T. Guillemaud, D. Haelewaters, A. Herz, A. Honek, A. G. Howe, C. Hui, W. D. Hutchison, M. Kenis, R. L. Koch, J. Kulfan, L. Lawson Handley, E. Lombaert, A. Loomans, J. Losey, A. O. Lukashuk, D. Maes, A. Magro, K. M. Murray, G. S. Martin, Z. Martinkova, I. A. Minnaar, O. Nedved, M. J. Orlova-Bienkowskaja, N. Osawa, W. Rabitsch, H. P. Ravn, G. Rondoni, S. L. Rorke, S. K. Ryndevich, M.-G. Saethre, J. J. Sloggett, A. O. Soares, R. Stals, M. C. Tinsley, A. Vandereycken, P. van Wielink, S. Viglášová, P. Zach, I. A. Zakharov, T. Zaviezo, and Z. Zhao. 2016. The harlequin ladybird, *Harmonia axyridis*: global perspectives on invasion history and ecology. Biological Invasions. 18: 997–1044.

Rudolf, V. H. W. 2019. The role of seasonal timing and phenological shifts for species coexistence. Ecology Letters. 22: 1324–1338.

Schaefer, P. W., R. J. Dysart, and H. B. Specht. 1987. North American distribution of *Coccinella septempunctata* (Coleoptera: Coccinellidae) and Its mass appearance in coastal Delaware. Environmental Entomology. 16: 368–373.

Schulte, L. A., B. E. Dale, S. Bozzetto, M. Liebman, G. M. Souza, N. Haddad, T. L. Richard, B. Basso, R. C. Brown, J. A. Hilbert, and J. G. Arbuckle. 2021. Meeting global challenges with regenerative agriculture producing food and energy. Nature Sustainability.

Seebens, H., T. M. Blackburn, E. E. Dyer, P. Genovesi, P. E. Hulme, J. M. Jeschke, S. Pagad, P. Pyšek, M. Winter, M. Arianoutsou, S. Bacher, B. Blasius, G. Brundu, C. Capinha, L. Celesti-Grapow, W. Dawson, S. Dullinger, N. Fuentes, H. Jäger, J. Kartesz, M. Kenis, H. Kreft, I. Kühn, B. Lenzner, A. Liebhold, A. Mosena, D. Moser, M. Nishino, D. Pearman, J. Pergl, W. Rabitsch, J. Rojas-Sandoval, A. Roques, S. Rorke, S. Rossinelli, H. E. Roy, R. Scalera, S. Schindler, K. Štajerová, B. Tokarska-Guzik, M. van Kleunen, K. Walker, P. Weigelt, T. Yamanaka, and F. Essl. 2017. No saturation in the accumulation of alien species worldwide. Nature Communications. 8: 14435.

Sethuraman, A., F. J. Janzen, M. A. Rubio, Y. Vasquez, and J. J. Obrycki. 2018. Demographic histories of three predatory lady beetles reveal complex patterns of diversity and population size change in the United States. Insect Science. 25: 1065–1079.

Shea, K. and P. Chesson. 2002. Community ecology theory as a framework for biological invasions. Trends in Ecology & Evolution. 17: 170–176.

Singer, M. C., and C. Parmesan. 2010. Phenological asynchrony between herbivorous insects and their hosts: signal of climate change or pre-existing adaptive strategy? Philosophical Transactions of the Royal Society B: Biological Sciences. 365: 3161–3176.

Sloggett, J. J. 2021. Aphidophagous ladybirds (Coleoptera: Coccinellidae) and climate change: a review. Insect Conservation and Diversity. 14: 709–722.

Smith, C. A., and M. M. Gardiner. 2013. Biodiversity loss following the introduction of exotic competitors: Does intraguild predation explain the decline of native lady beetles? PLOS ONE. 8: e84448.

Snyder, W. E. 2009. Coccinellids in diverse communities: Which niche fits? Biological Control. 51: 323–335.

Snyder, W. E., G. M. Clevenger, and S. D. Eigenbrode. 2004. Intraguild predation and successful invasion by introduced ladybird beetles. Oecologia. 140: 559–565.

Speights, C. J., and B. T. Barton. 2019. Timing is everything: Effects of day and night warming on predator functional traits. Food Webs. 21: e00130.

Thomas, A. P., J. Trotman, A. Wheatley, A. Aebi, R. Zindel, and P. M. J. Brown. 2013. Predation of native coccinellids by the invasive alien *Harmonia axyridis* (Coleoptera: Coccinellidae): Detection in Britain by PCR-based gut analysis. Insect Conservation and Diversity. 6: 20–27.

Turnock, W. J., and I. L. Wise. 2004. Density and survival of lady beetles (Coccinellidae) in overwintering sites in Manitoba. Canadian Field-Naturalist. 118: 309–317.

Vandereycken, A., D. Durieux, É. Joie, É. Haubruge, and F. Verheggen. 2012. Habitat diversity of the Multicolored Asian ladybeetle *Harmonia axyridis* Pallas (Coleoptera: Coccinellidae) in agricultural and arboreal ecosystems: a review. Biotechnologie, Agronomie, Societé et Environnement. 16: 553–563.

Vilà, M., J. L. Espinar, M. Hejda, P. E. Hulme, V. Jarošík, J. L. Maron, J. Pergl, U. Schaffner, Y. Sun, and P. Pyšek. 2011. Ecological impacts of invasive alien plants: a meta-analysis of their effects on species, communities and ecosystems. Ecology Letters. 14: 702–708. https://doi.org/10.1111/j.1461-0248.2011.01628.x

Vitousek, P. M., C. M. D’Antonio, L. L. Loope, and R. Westbrooks. 1996. Biological invasions as global environmental change. American Scientist.

Ware, R. L., and M. E. N. Majerus. 2008. Intraguild predation of immature stages of British and Japanese coccinellids by the invasive ladybird *Harmonia axyridis*, pp. 169–188. *In* Roy, H.E., Wajnberg, E. (eds.), From Biological Control to Invasion: The Ladybird Harmonia axyridis as a Model Species. Springer Netherlands, Dordrecht.

Wood, S. N. 2006. Generalized additive models: an introduction with R. chapman and hall/CRC.

Wu, Y., J. Li, H. Liu, G. Qiao, and X. Huang. 2020. Investigating the impact of climate warming on phenology of aphid pests in China using long-term historical data. Insects. 11.

Xue, Y., Bahlai, C.A., Frewin, A., Sears, M.K., Schaafsma, A.W. and Hallett, R.H., 2009. Predation by *Coccinella septempunctata* and *Harmonia axyridis* (Coleoptera: Coccinellidae) on *Aphis glycines* (Homoptera: Aphididae). Environmental Entomology, 38(3), pp.708–714.

Yasuda, H., E. W. Evans, Y. Kajita, K. Urakawa, and T. Takizawa. 2004. Asymmetric larval interactions between introduced and indigenous ladybirds in North America. Oecologia. 141: 722–731.

Yasuda, H., T. Kikuchi, P. Kindlmann, and S. Sato. 2001. Relationships between attack and escape rates, cannibalism, and intraguild predation in larvae of two predatory ladybirds. Journal of Insect Behavior. 14: 373–384.

Zaviezo, T., A. O. Soares, and A. A. Grez. 2019. Interspecific exploitative competition between *Harmonia axyridis* and other coccinellids is stronger than intraspecific competition. Biological Control. 131: 62–68.

