## Appendix S1 for "Coexistence between similar invaders: The case of two cosmopolitan exotic insects"

**Figure S1.** Pearson correlation results for environmental parameters used in within-season generalized additive model selection. Parameters are, from left to right, mean daily precipitation within a sampling week, number of rainy days within the sampling week, precipitation accumulation to date of sampling week, maximum daily rainfall observed within a sampling week, mean temperature during the sampling week, minimum temperature observed during the sampling week, maximum temperature observed during the sampling week, the degree day accumulation observed during the sampling week, and the degree day accumulation to date during that sampling week. Pearson correlation for each pair of parameters is given above the diagonal, a scatterplot of each combination of variables is given below the diagonal. The diagonal gives a frequency histogram of values for each different parameter.

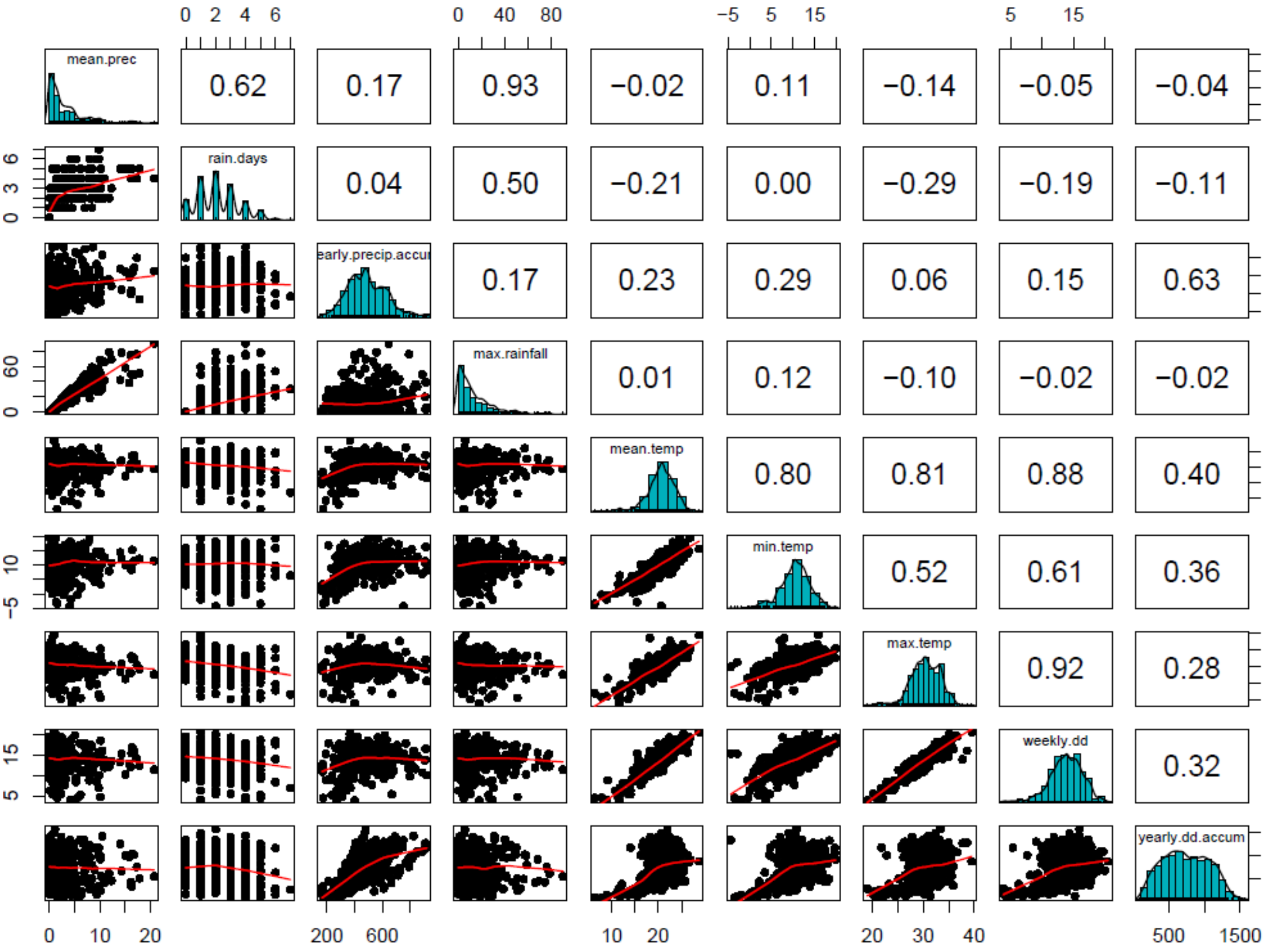

**Table S1.** Complete model selection results for within-year models explaining the spatio-temporal distribution of two lady beetle species from relative capture counts. Model specification used the sum of adult captures in a given plot, in a given observation period as the response variable, and included smoothed terms for environmental variables and year, and habitat as a parametric term. All terms interacted with species identity. Because of a high degree of collinearity between many predictor variables, a forward-selection approach was used to ensure no terms with Pearson correlation >0.5 appeared together in the model. An offset variable, the natural logarithm of the number of traps reporting for a given observation, was used to account for variation in sampling effort. All smoothing parameters were constrained to sp=1 to limit over-fitting. A quasi-Poisson error structure was used.

| Model | Habitat | Year | Precipitation term | Temperature term | Other terms | -REML | Adj $R^2$ | Deviance explained | Highest concurvity | Proportion of deviance from environmental terms | Notes |
|---|---|---|---|---|---|---|---|---|---|---|---|
| No environmental | X | | | | | 30520 | 0.07 | 14.9 | | | |
| | X | X | | | | 28897 | 0.13 | 25.0 | | | |
| Single environmental parameter | X | X | rainy days in week | | | 28836 | 0.14 | 25.4 | | 0.016 | Precipitation term constrained to 3 knots |
| | X | X | mean daily precipitation | | | 28622 | 0.17 | 26.8 | | 0.067 | |
| | X | X | max daily rainfall | | | 28627 | 0.16 | 26.8 | | 0.067 | |
| | X | X | precipitation accumulation (year to date) | | | 27848 | 0.21 | 31.1 | | 0.196 | |
| | X | X | | minimum temperature | | 28580 | 0.15 | 27.0 | | 0.074 | |
| | X | X | | mean temperature | | 28550 | 0.15 | 27.1 | | 0.077 | |
| | X | X | | maximum temperature | | 28466 | 0.16 | 27.6 | | 0.094 | |
| | X | X | | weekly degree day accumulation | | 28354 | 0.17 | 28.6 | | 0.126 | |
| | X | X | | degree day accumulation (year to date) | | 27426 | 0.22 | 33.1 | | 0.245 | |

| | | | | | | | | | | | |
|---|---|---|---|---|---|---|---|---|---|---|---|
| Best temperature parameter + uncorrelated precip | X | X | rainy days in week | degree day accumulation (year to date) | | 27394 | 0.226 | 33.3 | 0.212 | 0.249 | Precipitation term constrained to 3 knots |
| | X | X | mean daily precipitation | degree day accumulation (year to date) | | 27227 | 0.256 | 34.1 | 0.189 | 0.267 | |
| | X | X | max daily rainfall | degree day accumulation (year to date) | | 27152 | 0.256 | 34.7 | 0.197 | 0.230 | Best model with two environmental terms |
| Best precipitation parameter + uncorrelated temperature | X | X | precipitation accumulation (year to date) | minimum temperature | | 27385 | 0.244 | 33.8 | 0.496 | 0.260 | |
| | X | X | precipitation accumulation (year to date) | mean temperature | | 27287 | 0.251 | 34.3 | 0.471 | 0.271 | |
| | X | X | precipitation accumulation (year to date) | maximum temperature | | 27300 | 0.253 | 34.1 | 0.435 | 0.267 | |
| | X | X | precipitation accumulation (year to date) | weekly degree day accumulation | | 27211 | 0.264 | 34.8 | 0.432 | 0.282 | |
| Best temperature parameter + best uncorrelated precip + other uncorrelated params | X | X | max daily rainfall | degree day accumulation (year to date) | minimum temperature | 26927 | 0.275 | 36 | 0.606 | 0.306 | |
| | X | X | max daily rainfall | degree day accumulation (year to date) | mean temperature | 26936 | 0.269 | 35.9 | 0.648 | 0.304 | |
| | X | X | max daily rainfall | degree day accumulation (year to date) | maximum temperature | 26835 | 0.281 | 36.4 | 0.376 | 0.313 | Best overall model |

**Table S2.** Within-season multivariate model results explaining the spatio-temporal distribution of the two lady beetle species from relative capture counts (-REML=26835, n=27432, Deviance explained = 36.4%). Model specification used the sum of adult captures in a given plot, in a given observation period as the response variable, and included smoothed terms for degree day accumulation, maximum rainfall, maximum temperature, and year, and habitat as a parametric term. All terms interacted with species identity. An offset variable, the natural logarithm of the number of traps reporting for a given observation, was used to account for variation in sampling effort. All smoothing parameters were constrained to sp=1 to limit over-fitting. A quasi-Poisson error structure was used.

| Term | Effective degrees of freedom | F | p-value |
|---|---|---|---|
| Degree days : C7 | 6.26 | 169.96 | <0.001 |
| Degree days : HA | 6.76 | 92.74 | <0.001 |
| Max rainfall : C7 | 6.52 | 34.77 | <0.001 |
| Max rainfall : HA | 6.96 | 10.94 | <0.001 |
| Max temperature : C7 | 6.61 | 43.29 | <0.001 |
| Max temperature : HA | 6.71 | 15.01 | <0.001 |
| Year : C7 | 7.17 | 136.59 | <0.001 |
| Year : HA | 7.06 | 134.64 | <0.001 |
| Habitat (parametric term) | 8 | 165.14 | <0.001 |
| Species (parametric term) | 1 | 10.12 | <0.001 |
| Habitat x Species | 8 | 90.41 | <0.001 |

**Figure S2.** Pearson correlation results for environmental parameters used in between-season generalized additive model selection. Parameters are, from left to right, degree day accumulation at 20 weeks, 25 weeks. 30 weeks and 35 weeks, and then precipitation at 20 weeks, 25 weeks, 30 and 35 weeks of year. Pearson correlation for each pair of parameters is given above the diagonal, a scatterplot of each combination of variables is given below the diagonal. The diagonal gives a frequency histogram of values for each different parameter.

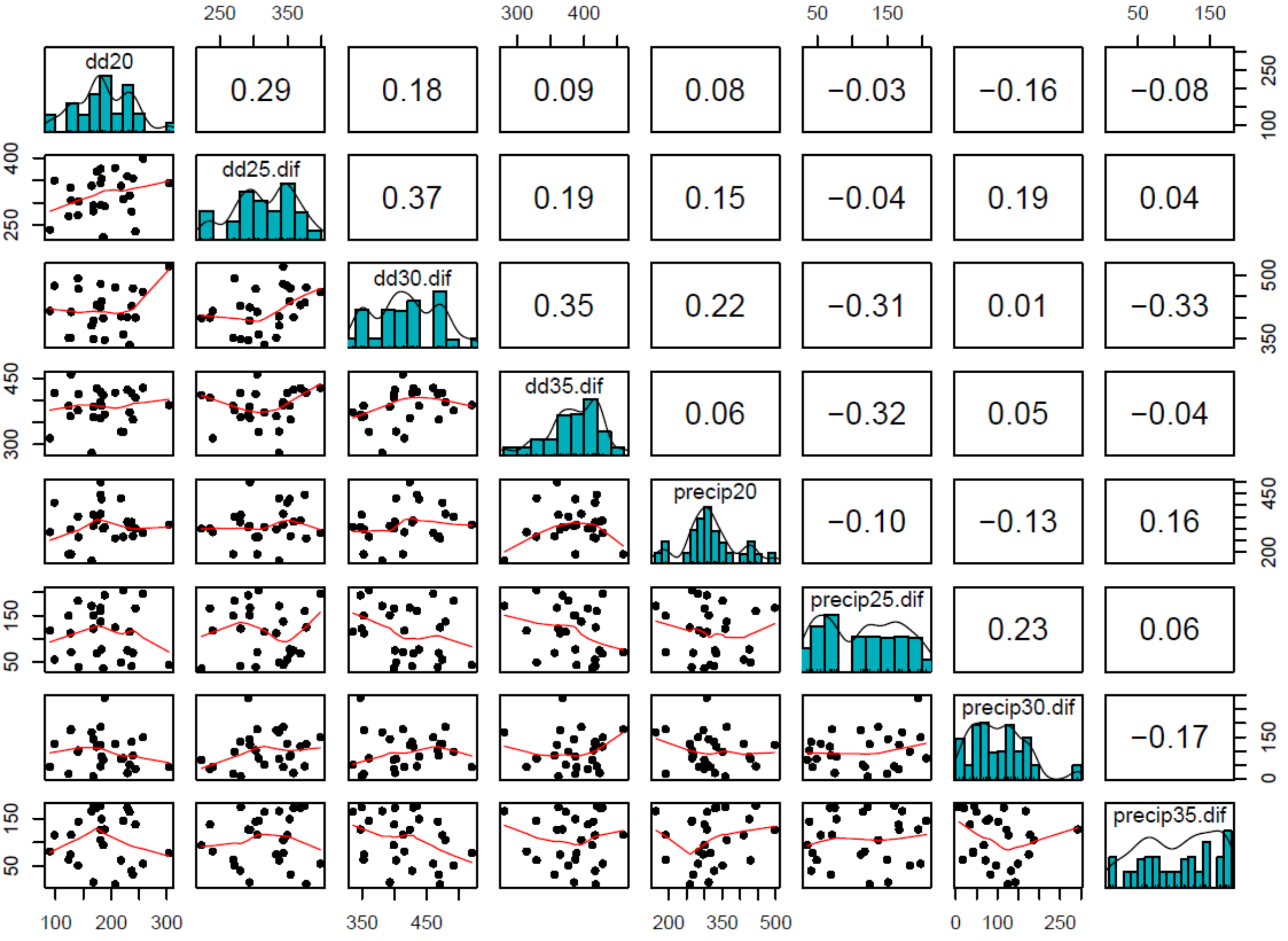

**Table S3.** Complete model selection results for between-year models explaining the spatio-temporal distribution of two lady beetle species from relative capture counts. Each species was modelled separately. Model specification used the sum of adult captures in a given plot, in a given observation period as the response variable, and environmental parameters were subjected to backward selection as described in the methods, with best model selected based on lowest -REML with no parameters with concurvity >0.8 . An offset variable, the natural logarithm of the number of traps reporting for a given observation, was used to account for variation in sampling effort. All smoothing parameters were constrained to sp=1 to limit over-fitting. A quasi-Poisson error structure was used. Variables in table are defined in Figure S2 caption.

| Species | Habitat | dd20 | dd25 | dd30 | dd35 | p20 | p25 | p30 | p35 | -REML | Adj R2 | Deviance explained | Highest concurvity (observed) | Proportion of deviance explained by environmental terms | Notes |
|---|---|---|---|---|---|---|---|---|---|---|---|---|---|---|---|
| HA | X | | | | | | | | | 2370 | 0.22 | 22.8 | | | |
| | X | X | X | X | X | X | X | X | X | 1844 | 0.67 | 68.2 | 1.000 | 0.666 | |
| | X | X | X | X | | X | X | X | X | 1842 | 0.67 | 68.2 | 1.000 | 0.666 | |
| | X | X | X | X | | X | | X | X | 1840 | 0.67 | 68.2 | 1.000 | 0.666 | |
| | X | X | X | X | | X | | | X | 1838 | 0.67 | 68.2 | 1.000 | 0.666 | Best HA model |
| | X | X | | X | | X | | | X | 1841 | 0.67 | 68.0 | 1.000 | 0.665 | |
| | X | | | X | | X | | | X | 1855 | 0.67 | 67.3 | 0.759 | 0.661 | Best HA model without concurvity issues |
| | X | | | | | X | | | X | 1967 | 0.59 | 59.9 | 0.489 | 0.619 | |
| C7 | X | | | | | | | | | 2552 | 0.20 | 36.6 | | | |
| | X | X | X | X | X | X | X | X | X | 1837 | 0.75 | 80.7 | 1.000 | 0.546 | |
| | X | X | X | X | X | X | X | X | | 1836 | 0.75 | 80.7 | 1.000 | 0.546 | |
| | X | | X | X | X | X | X | X | | 1838 | 0.75 | 80.6 | 1.000 | 0.546 | Best C7 model |
| | X | | X | X | X | | X | X | | 1846 | 0.75 | 80.3 | 1.000 | 0.544 | |
| | X | | X | | X | | X | X | | 1871 | 0.75 | 79.9 | 1.000 | 0.542 | |
| | X | | X | | X | | | X | | 1905 | 0.74 | 79.0 | 0.771 | 0.537 | Best C7 model without concurvity issues |
| | X | | X | | X | | | | | 2029 | 0.69 | 74.1 | 0.383 | 0.506 | |

**Table S4.** Between-year multivariate model results explaining the spatio-temporal distribution of the two lady beetle species from relative capture counts. The C7 model: -REML=1905.3, n=1353, Deviance explained = 79.0%; HA model: -REML=1855.1, n=1353, Deviance explained = 67.3%. Each species was modelled separately. Model specification used the sum of adult captures in a given plot, in a given observation period as the response variable, and environmental parameters were subjected to backward selection as described in the methods. An offset variable, the natural logarithm of the number of traps reporting for a given observation, was used to account for variation in sampling effort. All smoothing parameters were constrained to sp=1 to limit over-fitting. A quasi-Poisson error structure was used.

| Term | Effective degrees of freedom | F | p-value |
|---|---|---|---|
| C7 model | | | |
| Degree days at 25 weeks | 7.84 | 56.89 | <0.001 |
| Degree days at 35 weeks | 7.05 | 84.68 | <0.001 |
| Precipitation at 30 weeks | 5.14 | 43.08 | <0.001 |
| Habitat (parametric term) | 8 | 122.1 | <0.001 |
| HA model | | | |
| Degree days at 30 weeks | 6.02 | 26.06 | <0.001 |
| Precipitation at 20 weeks | 7.86 | 28.37 | <0.001 |
| Precipitation at 35 weeks | 6.56 | 41.59 | <0.001 |
| Habitat (parametric term) | 8 | 72.13 | <0.001 |
